# The Neural Determinants of Beauty

**DOI:** 10.1101/2021.05.21.444999

**Authors:** Taoxi Yang, Arusu Formuli, Marco Paolini, Semir Zeki

## Abstract

What are the conditions that determine whether the medial orbito-frontal cortex (mOFC), in which activity correlates with the experience of beauty derived from different sources, becomes co-active with sensory areas of the brain during the experience of sensory beauty? We addressed this question by studying the neural determinants of facial beauty. The perception of faces correlates with activity in a number of brain areas, but only when a face is perceived as beautiful is the mOFC also engaged. The enquiry thus revolved around the question of whether a particular pattern of activity, within or between areas implicated in face perception, emerges when a face is perceived as beautiful, and which determines that there is, as a correlate, activity in mOFC. 17 subjects of both genders viewed and rated facial stimuli according to how beautiful they perceived them to be while the activity in their brains was imaged with functional magnetic resonance imaging. A univariate analysis revealed parametrically scaled activity within several areas in which the strength of activity correlated with the declared intensity of the aesthetic experience of faces; the list included the mOFC and two core areas strongly implicated in the perception of faces - the occipital face area (OFA), fusiform face area (FFA)- and, additionally, the cuneus. Multivariate analyses showed strong patterns of activation in the FFA and the cuneus and weaker patterns in the OFA and the pSTS. It is only when specific patterns emerged in these areas that there was co-activation of the mOFC, in which a strong pattern also emerged during the experience of facial beauty. A psychophysiological interaction analysis with mOFC as the seed area revealed the involvement of the right FFA and the right OFA. We conjecture that these collective patterns of activity constitute the neural basis for the experience of facial beauty, bringing us a step closer to understanding the neural determinants of aesthetic experience.

## Main

The work reported here represents the beginnings of an exploration of what determines the beauty of objects. In the age-old discussions about the determinants of beauty, the brain has not played an explicit role, assuming it to have played one at all. Nevertheless, the notion that beauty’s determinants may also lie in the brain, rather than solely in the external world, has been hinted at sporadically, but usually only implicitly and vaguely. Although Plato himself hardly ever mentioned the brain1, knowledge of beauty in his Theory of Forms can only be aspired to, but never completely attained, by a thought process, implying an involvement of the brain; the writings and recorded thoughts of other philosophers and artists, among them the Greek sculptor Polykleitos, the Alexandrian Greek neo-Platonist philosopher Plotinus, as well as Michelangelo and Leonardo, similarly hint implicitly at the involvement of thought processes, and therefore of the brain, in determining beauty (Clements, 1961; Nicholl, 2004; Tobin 1975; Plotinus, 1964). The critical involvement of the brain was explicitly acknowledged only with the demonstration that the experience of beauty, regardless of its source, correlates with quantifiable activity in a specific part of the emotional brain, namely the mOFC (Blood et al., 2001; Ishizu & Zeki 2011; Kawabata & Zeki 2004; Tsukiura & Cabezza, 2011; Zeki et al., 2014, inter alia); that demonstration raises the important but unaddressed issue of learning what objective characteristics of apprehended objects leads to activity in the sensory areas of the brain which have, as a consequence, the engagement of the mOFC as well, for only when a sensory stimulus is experienced as beautiful is the mOFC engaged along with the sensory areas.

To address this question experimentally, we restricted ourselves to studying one category of beautiful stimuli, namely that falling into the biological category (Zeki & Chen, 2020) and, within that category, confined ourselves further to studying the neurobiology underlying the experience of facial beauty. Even such a restriction presents considerable difficulties because, in spite of a general agreement about the aesthetic status of a face when it is very beautiful (Bignardi et al., 2020; Vessel et al., 2019), there is no consensus about what critical “objective” feature constitutes the basis for this agreement. Among the features considered necessary for rendering a face beautiful are certain proportions and symmetries, in addition to what some have considered to be exact, mathematically defined, relations between parts, as enshrined in particular in the golden ration; but there has been no unanimity of views on this (Alam et al., 2015; Cellucci, 2017; Hönn & Göz, 2007; Jones & Hill 1993; Swift & Remington, 2011). What is certain is that there is another elusive and ineffable quality, referred to by Clive Bell as a “mysterious law” (Bell, 1914) and which, in an imaginary dialogue, Hofstadter (1980) summarizes as follows: “Indeed, I would venture to say there exists no set of rules which delineate what it is that makes a piece beautiful, nor could there ever exist such a set of rules. The sense of Beauty is the exclusive domain of Conscious Minds, minds which through the experience of living have gained a depth that transcends explanation by any mere set of rules”. Could that extra, mysterious, ingredient, be a certain “significant configuration” (Zeki, 2013) that reveals itself as a distinct pattern of activity in specific face processing areas of the brain, a pattern that may constitute the necessary neural ingredient that renders a face beautiful. That is the cardinal question addressed here.

### Brain areas implicated in face perception

The viewing of faces leads to activity in several areas of the brain. The first to be described and the one that has received most attention is the fusiform face area (FFA) (Kanwisher & Yovel 2006; Sergent et al., 1992). Yet there are many other areas that play different roles in face perception; among these is the occipital face area (OFA) (Allison et al., 1999; Gauthier et al., 2000; Pitcher et al., 2007, 2011) and an area located in the superior temporal sulcus (pSTS) (Phillips et al., 1997; Puce at al., 1998) involved in registering facial expressions (Engell & Haxby, 2007), as well as other areas, detailed in the Discussion, whose function may not be specific or limited to face perception. Hence, the current consensus is that there is a network of distributed areas that are involved in various aspects of face processing (Haxby et al., 2000; Ishai, 2007; Ishai et al., 2005; Kanwisher & Barton 2011). But, while the perception of faces results in activity within a number of cortical areas, only if the face is experienced as beautiful is there, in addition, activity in the mOFC (Bartels & Zeki 2004; Ishai, 2007; O’Doherty et al., 2003; Winston et al., 2007). This implies that a selection occurs, according to criteria yet to be established, in one or more of the cortical areas that are responsive to faces, which leads to co-activation of mOFC, either directly or indirectly. Our specific hypothesis was that, whatever their precise details, objective characteristics of stimuli that result in their being experienced as beautiful do so because they trigger particular patterns of activity within one or more sensory areas critical for face perception; it is only when such specific patterns emerge in the sensory (face) areas that there is co-activity in the mOFC, with the experience of beauty as a correlate. By limiting ourselves to the experience of facial beauty, we have tried to address, in an experimentally more manageable though limited way, the general question of what the neural determinants of beauty may be.

To do so, we extended earlier studies that have determined an apparently critical role for the mOFC during the experience of beauty, by studying the relationship between mOFC activity and that in different brain regions that are active when subjects experience (facial) beauty. Given that brain areas such as the FFA or the OFA, as well as other areas implicated in the perception of faces, respond to both average and beautiful faces as well as to ugly ones, it becomes important to learn whether their response to different aesthetic categories of faces differs in any way, for example in the strength or pattern of activity that correlates with a given aesthetic evaluation of a face. Such an approach is made possible by advances in multivariate pattern analyses (MVPA) (Haxby et al., 2001, 2014),including multivariate pattern classification (MVPC) and representational similarity analysis (RSA) (Kriegeskorte et al., 2008; Kriegeskorte, & Kievit, 2013).

Addressing this question carried with it the hope of gaining germs of an answer to another, and more general, neurobiological problem. That only under certain conditions (in this case aesthetic experience) does activity in the mOFC correlate with activity in the relevant sensory areas of the cortex implies that only when a selection occurs at some sensory level are the results of the processings in the sensory areas relayed, directly or indirectly, to the mOFC. The issue of selection is of general importance in cortical neurobiology: each cortical area has multiple outputs that differ in their destinations; this raises the question whether all these outputs are engaged when a given cortical area undertakes one of its operations, or whether they are engaged on a “need to know” basis only, since any given cortical area may undertake multiple tasks related to its specialization (Zeki, 2015).

The overall question that we address here can thus be summarised as follows: do certain, as yet undefined, objective characteristics of a facial stimulus result in particular patterns of activity within sensory perceptive areas for faces, patterns that can be reasonably supposed to lead to the experience of the perceived faces as beautiful because they also result in the co-activation of the mOFC? The question can also be phrased alternatively, by asking whether there any detectable neural configurations that emerge in face perceptive areas only during the experience of beautiful faces, and that these configurations are indicative of the presence of certain, as yet undefined, objective qualities in the stimulus. From this, it would follow that faces that are perceived to be neutral or ugly do not result in similar configurations because they lack those objective qualities.

That there are alternative ways of phrasing the question is testament to the age-old tension in the debate about beauty and whether it is in the object alone, the perceiver alone, or in both.

To answer these questions of how facial beauty is represented in the human brain, we presented 120 unique faces to 17 participants during fMRI. After each picture presentation (2 s), participants rated each face’s beauty level on 7-piont Likert scale (1–7, least to most) (Figure 1). our strategy was to ask subjects of both sexes and of different cultural and racial backgrounds to view pictures of faces in the scanner and rate them according to how beautiful they found them to be while the activity in their brains was being scanned. We tried to learn whether the experience of facial beauty can be decoded based on the activity patterns produced in in one or more areas when subjects experience different categories of facial beauty.

**Figure 1.**
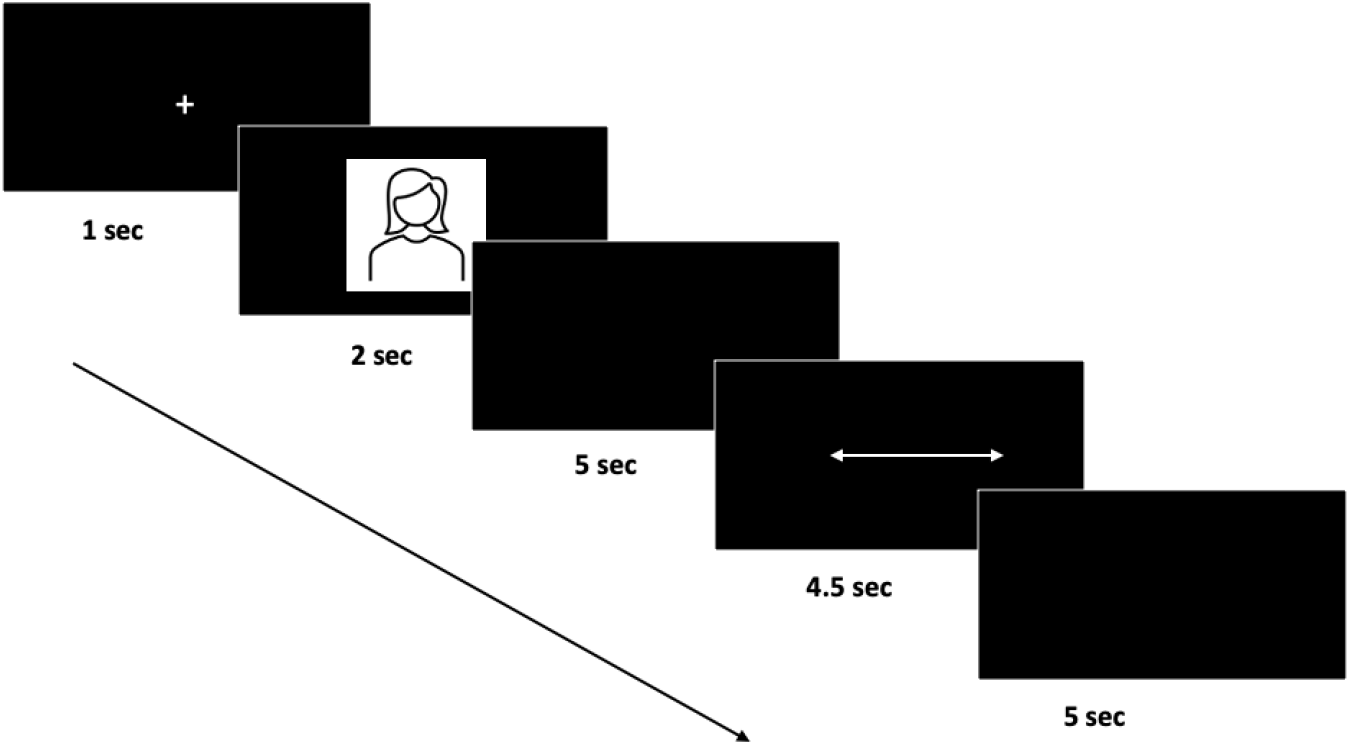
The sequence of presentation and response in the fMRI scanner, indicated by the direction of the arrow. Each trial began with a fixation cross (1 s) on a blank screen, followed by an image of a face (2 s) and a blank screen (5s). This was followed by a 4.5 s rating period during which the subject used buttons to indicate their response on a visual slider. After another 5-s blank screen, the next trial began.

## Results

### Behavioural Responses

A repeated-measures one-way analysis of variance (ANOVA) with 3 levels (“very beautiful”, “average”, and “not-beautiful”) was conducted on the beauty ratings that were obtained in the scanner. This showed that there was a significant difference in beauty ratings as a function of stimulus (F_2,78_ = 464.38, *p* < 0.0005, η^2^= 0.92). Follow-up repeated-measures t-tests showed that mean ratings were higher for the “very-beautiful” faces (mean = 5.80) compared to the “average” (mean = 3.51, p < 0.0005) and “not beautiful” ones (mean = 2.38, *p* < 0.0005). The difference in beauty ratings between “average” and “not beautiful” faces was also significant (*p* < 0.0005). Ratings of attractiveness after the scanning session are given in Supplementary materials. Faces in the “very beautiful” category were rated as being significantly more attractive than those in the “average” and “not beautiful” categories. The average attractiveness ratings in the three categories were 5.34, 2.86 and 1.93, respectively.

A comparison of beauty ratings with attractiveness ratings showed that there was a significant but not complete overlap between the two (Figure 2A). In particular, the overlap was lowest for faces rated low on the beauty scale and did not exceed 75% for faces given ratings of 4 (“average”) and 6 (“very beautiful”). Also shown in Figure 2B are the RDMs for the beauty and attractiveness ratings, from which it emerges that there is a higher agreement among subjects for rating beautiful faces as attractive and a considerable spread in agreement on the attractiveness of faces rated as “average” and “not beautiful”.

**Figure 2.**
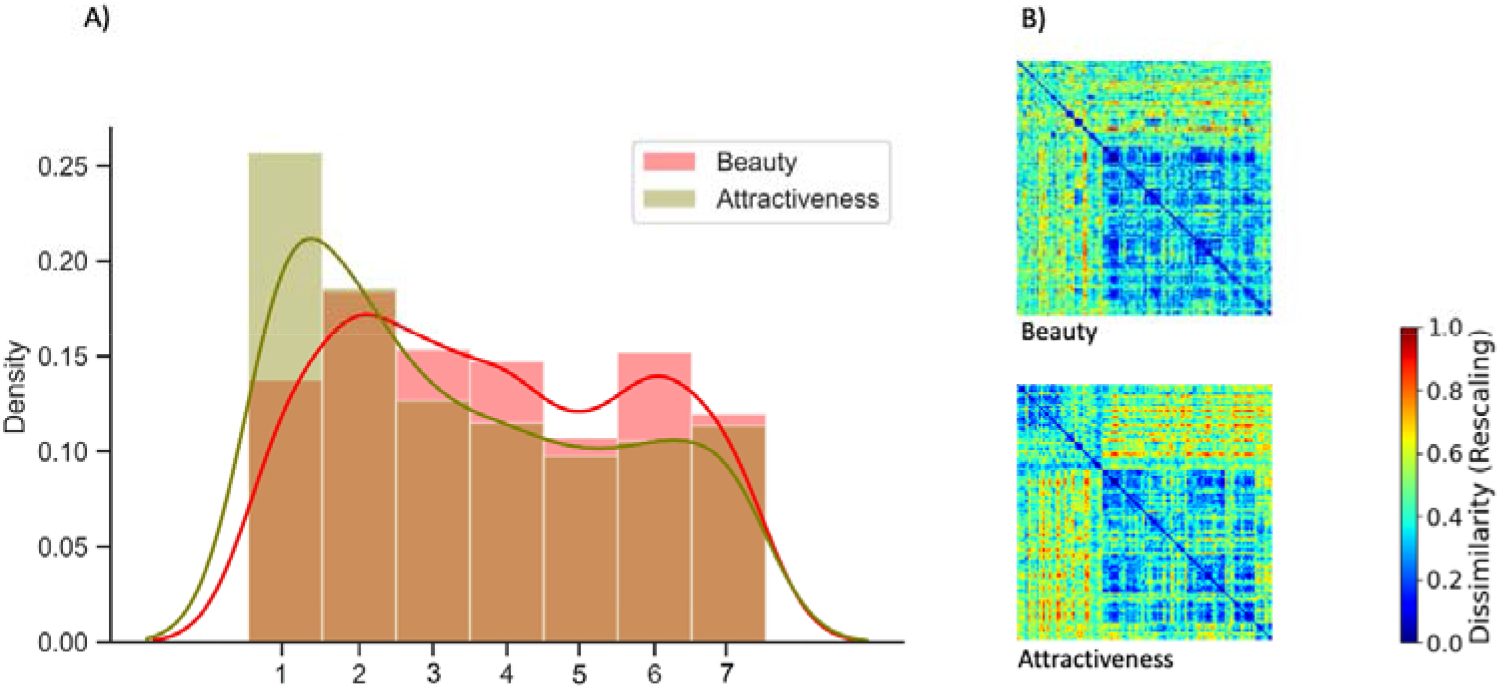
Comparison of the beauty ratings in relation to the attractiveness ratings in this study. A) The histogram and kernel smooth density plot for beauty (red) and attractiveness (green) ratings in all participants. B) Representational dissimilarity matrices (RDMs) for the beauty and attractiveness ratings. The RDMs represent pairwise comparisons between the 120 stimuli with regard to each of the beauty/attractiveness ratings. The dissimilarity measure reflects Spearman correlation, with blue indicating strong similarity and red strong dissimilarity. The RDMs were organised by average ratings decreasingly.

### Univariate analyses

First, a random effects analysis was applied to the contrast “very beautiful” vs not “not beautiful” faces, followed by a random effects analysis to search for regions in which activity was parametrically modulated by the participants’ beauty ratings.

#### Categorical analysis

The contrast “*very-beautiful*” > “*not-beautiful*” for faces revealed significant activations bilaterally in the fusiform face area (FFA) and the right occipital face area (OFA); additionally, there were significant clusters bilaterally in the cuneus and unilaterally in the left precuneus, as well as in the medial orbital frontal cortex (mOFC). No significant clusters were observed for the contrast “*not-beautiful”* > “*very-beautiful*” (See Table 1). Figure 3A shows the resulting statistical maps.

**Table 1.** Location of active brain regions in the comparison of “very-beautiful” vs “not-beautiful” face conditions. The reverse comparison (“not-beautiful” vs. “very-beautiful”) did not reach significance. Regions are designated using the MNI coordinates. L indicates left hemisphere, and R indicates right hemisphere. All results thresholded at p < 0.0001 at voxel level and FWE corrected p < 0.05 at cluster level.

| Brain regions | MNI coordinates |  |  | Max *t* scores | Number of voxels |
| --- | --- | --- | --- | --- | --- |
|  | x | y | z |  |  |
| L Cuneus | -30 | -85 | 23 | 7.20 | 154 |
| R Cuneus | 21 | -88 | 23 | 6.10 | 173 |
| L Precuneus | -3 | -55 | 14 | 5.55 | 28 |
| L FFA | -30 | -67 | -16 | 5.15 | 63 |
| R FFA | 30 | -62 | -16 | 4.56 | 81 |
| R OFA | 30 | -70 | -16 | 4.33 |  |
| L Medial Orbital Frontal Cortex | -3 | 53 | -4 | 4.39 | 100 |

**Figure 3.**
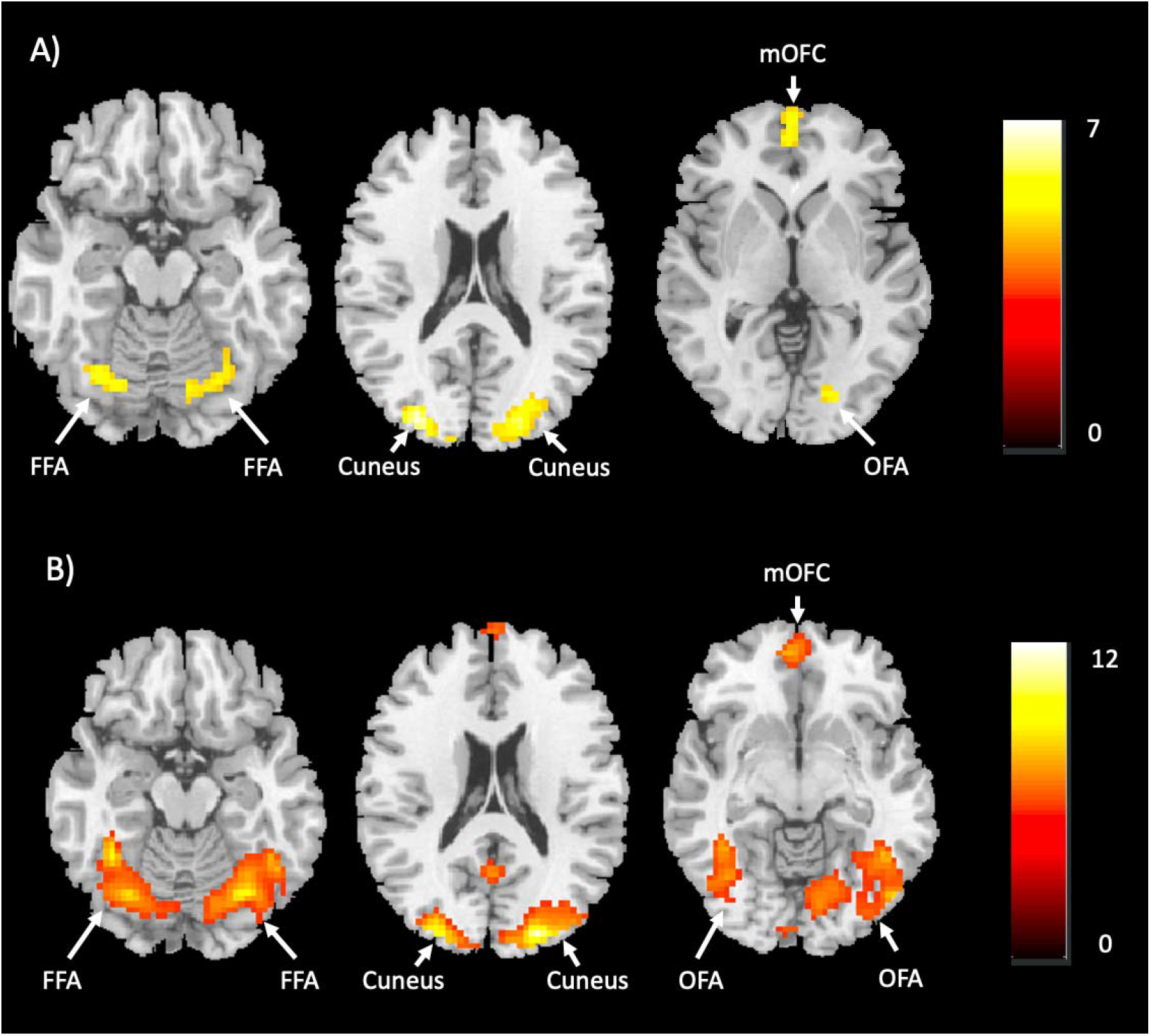
A) Categorical analysis results. The “very-beautiful” vs the “not-beautiful” faces contrast revealed activations bilaterally in the fusiform face area (FFA) and the cuneus, the right occipital face area (OFA) and the bilateral medial orbital frontal cortex (mOFC). Note. The left precuneus (not shown) was also active. B) Parametric analysis results, which revealed that the FFA and OFA, cuneus, and mOFC were active bilaterally. Note. Other activated areas included the left precenues, the right middle occipital gyrus, the right precentral gyrus, and the left superior frontal gyrus.

#### Parametric analysis

At the whole-brain level, we aimed to identify brain regions that showed a linear change in activity with the declared intensity of the experience of beauty in faces; the entire set of beauty rating for each face (1- 7) was used as the parametric regressor. Results showed activity in several brain regions (Table 2 & Figure 3B).

**Table 2.** Whole-brain parametric analysis of increasing beauty ratings. There was no significant activity correlated with decreasing beauty ratings. Regions are designated using the MNI coordinates. Conventions and thresholding as for Table 1.

| Brain regions | MNI coordinates |  |  | Max *t* scores | Number of voxels |
| --- | --- | --- | --- | --- | --- |
|  | x | y | z |  |  |
| R Middle Occipital Gyrus | 45 | -76 | 11 | 12.48 | 2059 |
| R Cuneus | 21 | -88 | 17 | 11.10 |  |
| R Occipital Face Area (OFA) | 48 | -73 | -7 | 7.35 |  |
| L Cuneus | -9 | -94 | 8 | 6.67 |  |
| R Fusiform Face Area (FFA) | 36 | -67 | -13 | 6.10 |  |
| L Fusiform Face Area (FFA) | -39 | -55 | -13 | 10.43 | 466 |
| L Occipital Face Area (OFA) | -39 | -76 | -4 | 5.08 |  |
| L Superior Frontal Gyrus | -18 | 38 | 47 | 9.43 | 57 |
| L Medial Orbital Frontal Cortex | -3 | 56 | -1 | 7.95 | 229 |
| R Precentral Gyrus | 45 | -7 | 53 | 7.74 | 42 |
| L Precuneus | -3 | -52 | 14 | 7.52 | 73 |

To exclude possible confounding factors that may result from familiarity, a parametric regressor representing familiarity was added to the design matrix used in the fMRI data analysis; no voxels showed significant activation associated with increasing familiarity ratings at the p⍰<⍰0.05, FWE corrected level. Thus, the differences observed in brain activations cannot be accounted for by familiarity.

#### Psychophysiological interactions (PPI) analysis

Given that the mOFC was found to play an important role in the experience of facial beauty (see above results), a further question that followed was: which brain regions does mOFC interact during the experience of beauty? A psychophysiological interaction analysis is computed to estimate condition-related changes in functional connectivity between brain areas (Friston et al., 1997; O’Reilly et al., 2012). One-sample t-test of the PPI analysis showed that, during the “very-beautiful” compared to the “not-beautiful” face conditions, the right OFA and right FFA displayed significantly increased connectivity with the mOFC (p⍰<⍰0.001) (see Table 3).

**Table 3.** Results from PPI analysis during high-beauty compared with low-beauty face conditions Note. Regions are designated using the MNI coordinates. R indicates right hemisphere. All results thresholded at p < 0.001 at voxel level and FWE corrected p < 0.05 at cluster level.

| Brain regions | MNI coordinates |  |  | Max *t* scores | Number of voxels |
| --- | --- | --- | --- | --- | --- |
|  | x | y | z |  |  |
| R OFA | 42 | -70 | -10 | 5.15 | 117 |
| R FFA | 48 | -58 | -13 | 4.57 |  |

### Multivariate analyses

MVPC analysis, based on a logistic regression classification algorithm (Hastie at al., 2009), was used to generate and evaluate how effective a cross-participant classifier would be in distinguishing the activity produced by viewing faces that fell into the “very-beautiful” and the “not-beautiful” categories. This decoding approach aims at predicting the class of stimuli viewed (“classification problem”) based on the pattern of brain activation elicited by them. The models are fitted on part of the data (train set) and tested on left–out data (test set). We used a cross-validation scheme that shuffles the subjects between the training and test sets, i.e., leave-one-subject-out. In the second RSA approach, we examined whether the representational geometry (the pattern of distance between voxels) in a given area, produced by the experience of “very-beautiful” faces, is different from that produced in the same area by the experience of “not beautiful” faces, in high-dimensional neural space. Our goal was to identify brain regions in which there may be fine-grained beauty-specific activity patterns, i.e., to learn whether the experience of facial beauty leads to a particular pattern of activity, on the basis of which the experience of beauty can be decoded when subjects experience different categories of facial beauty. Combining decoding analysis and RSA, we found that “very-beautiful” faces appear to be primarily encoded in face-selective areas (FFA, OFA and pSTS) as well as in the mOFC.

#### Decoding analysis

Results from the MVPC corresponded broadly with that derived from the univariate analysis, but the two maps did not overlap completely. Whole-brain “searchlight” analyses identified face-selective areas and the mOFC as containing information relevant for distinguishing between activations produced by the viewing of “very-beautiful” and “not-beautiful” faces. Significant classification was observed in the bilateral cuneus, the right OFA, the right FFA, the pSTS, all of them areas previously implicated in one way or another with the perception of faces (see Discussion). There was also bilateral activity in mOFC, already implicated in the experience of beauty derived from different sources, including faces (see Discussion). We investigated these regions further with analyses using probabilistic ROI masks; we found that the cross-participant classification of brain activity related to the experience of facial beauty across classes was highest in the mOFC (62.35%), followed by the right FFA (61.17%). The decoding accuracies of OFA, pSTS and cuneus were relatively lower but exceeded chance (53.53%, 55.29% and 57.06%, respectively) (Figure 4). Such cross-participant commonality may allow a common scaling of the experience of facial beauty across individuals.

**Figure 4.**
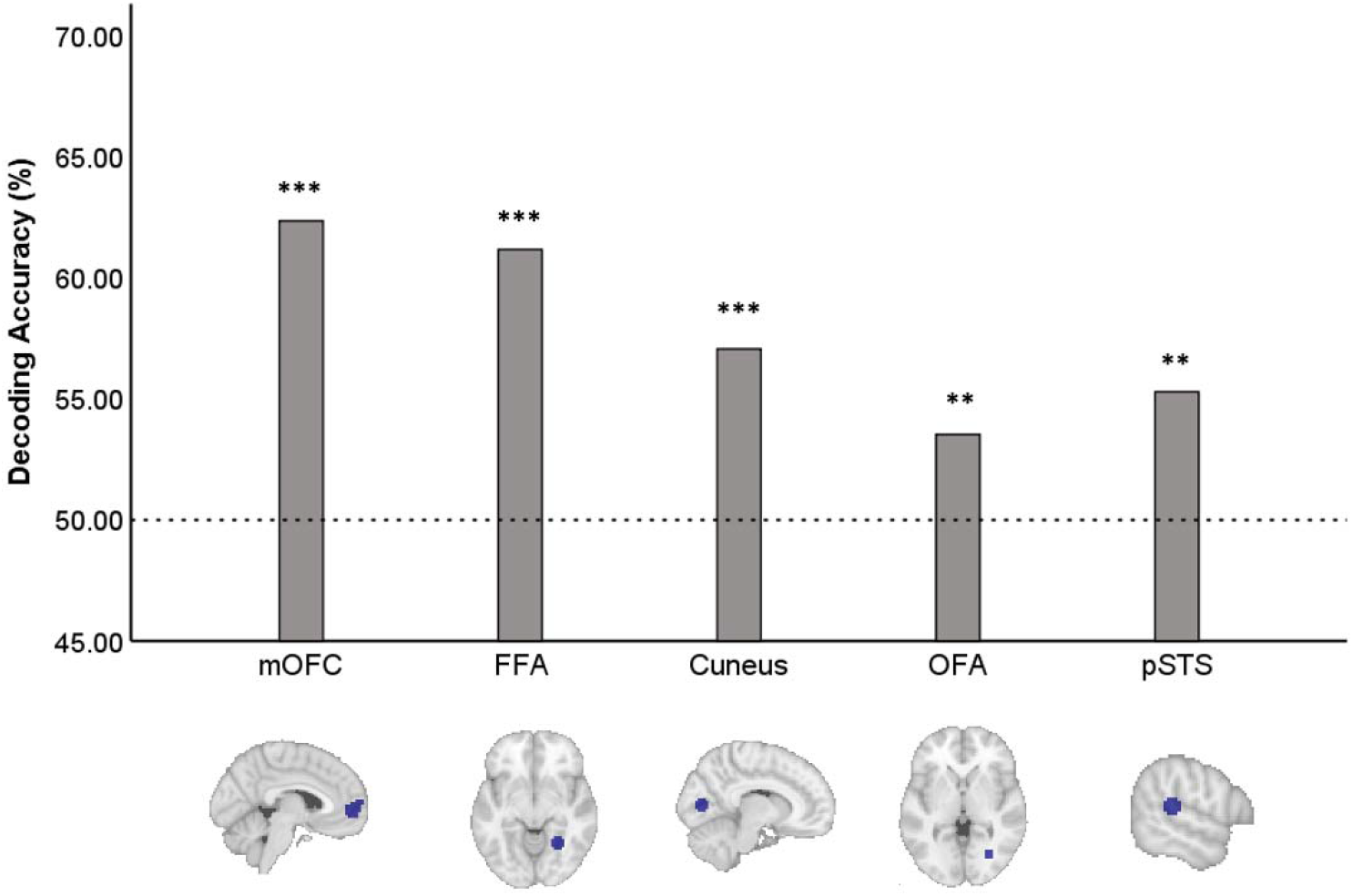
The cross-participant multi-voxel pattern classification (MVPC) accuracies within the individual ROIs. The dotted line signifies chance level accuracy (50%). \*\**p* < 0.01; \*\*\**p* < 0.001.

In sum, these findings suggest that, across people from different cultural and racial groups, the experience of facial beauty results in, or is based on, a common “code” in specific areas of the brain implicated in the perception of faces, as well as in the mOFC.

#### Representational similarity analysis

Two RSA models were examined; the first tested for responses in which the similarity was driven by the viewing of “very beautiful” faces. This model excluded the possibility that the similarity could be driven by responses to “not-beautiful” faces, by hypothesising that the representational patterns are not similar among “not-beautiful” faces (Figure 5C, Model 1). Using a whole-brain searchlight analysis, we identified several locations in which pattern similarity was sensitive to “very beautiful” faces. Among them are the left FFA, the right FFA, the right OFA, the right pSTS, and bilaterally in the cuneus and the mOFC.

**Figure 5.**
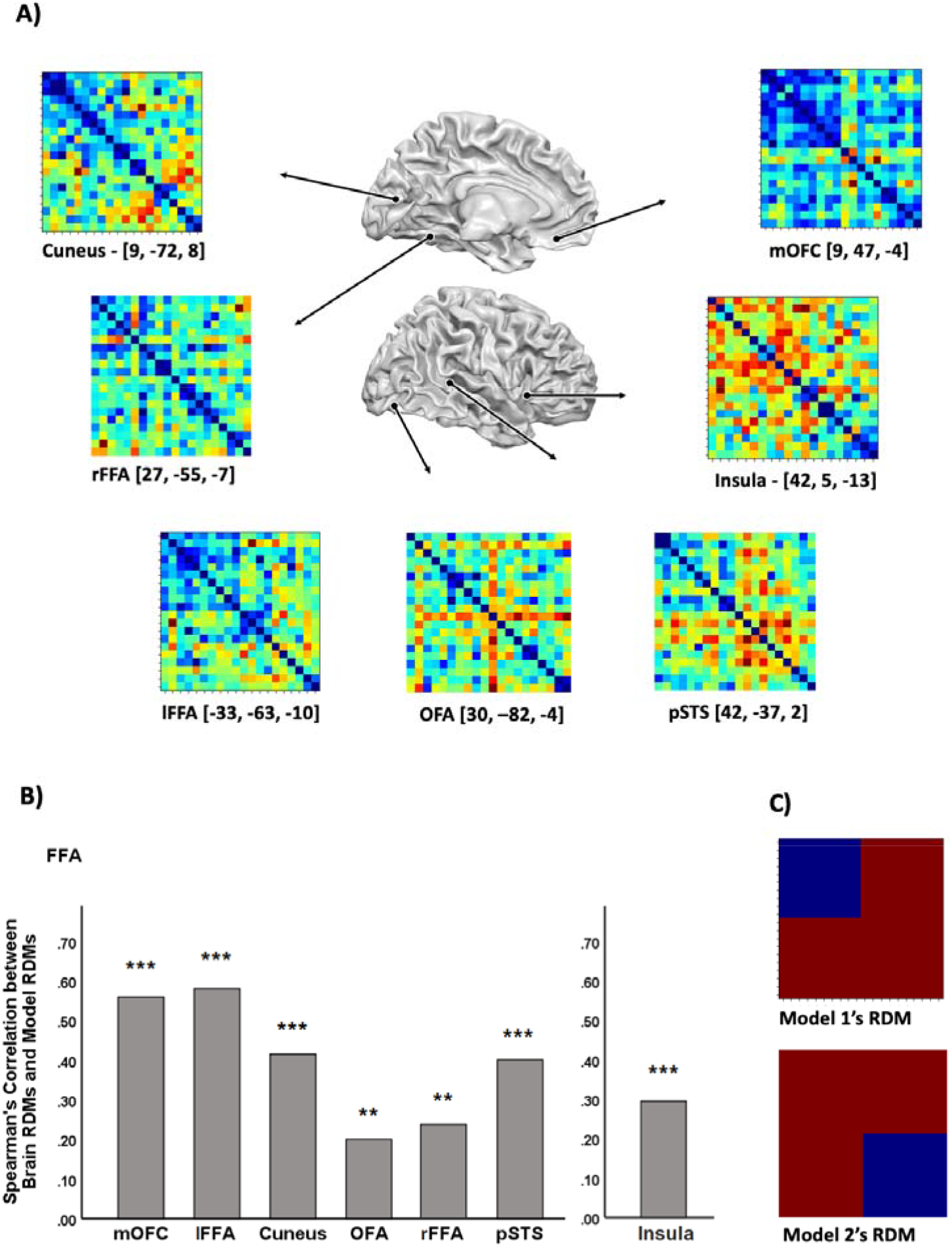
Representational similarity analysis (RSA) results. A) Brain RDMs averaged across subjects in the bilateral mOFC, bilateral FFA, bilateral cuneus, right OFA and right pSTS. For visualization purposes, distance metrics are scaled from 0 (most similar) to 1 (most dissimilar). B) Spearman corelation coefficients between brain and model representational dissimilarity matrices (RDMs). The left graph corresponds to model 1 and the right to model 2. \*\**p* < 0.001; \*\*\**p* < 0.0001. C) Two Model RDMs used for RSA, which indicates for each pair of faces whether they have the same or different values. Model 1 predicts that “very beautiful” faces evoke consistent patterns, while Model 2 predicts that “not beautiful” faces evoke consistent patterns.

The second model examined responses in which the similarity was driven by “not-beautiful” faces. This model tested for regions in which the response to “not-beautiful” faces showed similar patterns, irrespective of the patterns produced by the viewing of “very beautiful” faces (Figure 5C, Model 2). A whole-brain searchlight analysis revealed clusters of spotlight locations in the right insula, in which pattern similarity was greater for perceiving low-beauty faces.

Neural RDMs showing the discriminability of brain response patterns were created for each ROI (see above). Comparison of the neural RDM with the model RDMs in these ROIs allowed us to compare the manner in which “very-beautiful” and “not-beautiful” faces is represented in specific brain regions, enumerated above. The Spearman correlation coefficients between model RDMs and brain RDMs were as follows: mOFC, *r* = 0.56, p < 0.0001; left FFA, *r* = 0.58, *p* < 0.0001; right FFA, *r* = 0.23, *p* < 0.001; right OFA, *r* = 0.21, *p* < 0.001; right pSTS, *r* = 0.40, *p* < 0.0001; cuneus, *r* = 0.42, p < 0.0001 (see Figure 5B). This shows that the shared variance in the behavioural categorization of “very-beautiful” and the “not-beautiful” faces is also reflected in brain representational geometries, further confirming that these brain regions encode the categorical representations (aesthetic status) of the face stimuli, and that there is a significant neural representational geometry in the cortical areas implicated in face perception and in the mOFC for beautiful faces.

## Discussion

A “significant configuration” leading to aesthetic experience revealed in a neural pattern within sensory areas?

We set out to determine whether particular and detectable neural patterns emerge in one or more “face-perceptive” areas when humans experience a face as beautiful, our conjecture being that the emergence of such particular patterns constitute the condition for the engagement of mOFC and that only parallel activity in the “face perceptive” areas and mOFC lead to the experience of beauty. The results of the multivariate analyses reported here support our general hypothesis that significant classification patterns emerge in face perceptive areas during the experience of facial beauty: the areas implicated are the FFA, OFA, cuneus and pSTS, the first three being ones in which activity was also parametrically scaled with the declared intensity of the aesthetic experience. Among these areas, the highest cross-participant classification accuracy was in the right FFA and, while it remained above chance, the accuracy was lower in the other areas. The activity in these areas also correlated with activity in the mOFC, where the strongest representational similarity was found (see also Pegors et al., 2015). Taken together with similar results obtained from the multivariate RSA analysis, it seems reasonable to conclude that there is, within some of the areas implicated in face perception but especially in the FFA, OFA, cuneus and pSTS, a shared geometric representational similarity across subjects belonging to different ethnic and cultural groups which registers the aesthetic status of a face. Of these, the cuneus, unlike the other areas, has not occupied as prominent a position in the literature on face perception, although it has been sporadically charted (see Rossion et al., 2012); it has in particular been implicated in activities related to “theory of mind” and lip reading (Campbell et al., 2001; Gobbini & Haxby, 2007).

The areas listed above constitute a limited set from the more extensive set of areas in which activity is modulated by the declared intensity of the aesthetic experience of faces, which our parametric analysis revealed. Of these, the right middle occipital gyrus, like the right precentral gyrus, has been previously implicated in recording the emotional and rewarding properties of faces, among other stimuli (Dolan et al., 1966; Nakamura et al., 1998; Vartanian et al., 2013); the left superior temporal gyrus has been found to be involved in evaluating the emotional expression on faces (Del Casale et al., 2017; Kano et al. 2003; Kilts et al., 2003; Scheuerecker et al., 2007); the right precentral gyrus has been considered to be involved in empathetic facial processing and, more generally, in empathetic social cognition (Adolphs et al. 2000; Seehausen et al. 2016), while the superior frontal gyrus has been implicated in the perception of “talking” faces (Hall et al., 2005). That no particular representational similarity pattern emerged in these areas during the experience of facial beauty does not necessarily mean that they are not involved in unknown ways in rendering a face beautiful to the perceiver; they could have been involved, perhaps indirectly, by modulating activity in the areas in which a classification pattern emerged (e.g., the FFA, OFA and). It is therefore possible, and even likely, that the latter areas do not act in isolation but that the emergence of a pattern in them is dependent on inputs to them from other areas, particularly ones which our parametric analysis revealed.

It is trite, but nevertheless important, to emphasize that many features, both “static” and “dynamic”, taken together, render a face beautiful and that symmetry, proportion, and relationship of parts, while being critical, are not the sole ones. Viewed in this light, our finding, that the aesthetic experience of faces leads to widespread cortical activity, is not surprising; it reflects the wide distribution of cortical areas engaged in the perception of faces (Haxby et al., 2000; Ishai, 2005; Kanwisher & Barton 2011), with different areas emphasizing different facial features. But our PPI analysis revealed that, of the many areas active in our categorical and parametric analyses, only two - the right OFA and the right FFA - were in direct interaction with the mOFC. We are uncertain as to how much emphasis to place on the fact that activity in these two areas was unilateral. There have been earlier descriptions of lateralized activity in both areas, for example during early face processing (Pitcher et al., 2007) or during the determination of “self-face” identity (Ma & Han, 2011); this suggests that there may be a greater degree of specialization within areas that are critical for face perception.

### The anatomical connections between the active “face-perceptive” areas and the mOFC

Direct or indirect connections from the fusiform gyrus to the mOFC have been posited (Elbich et al., 2019; Fairhall & Ishai, 2007), and this includes the two areas, the right FFA and the right OFA, which our PPI analysis revealed. To our knowledge direct outputs from the other areas implicated in face perception revealed in our study have not been charted, which suggests that the FFA and the OFA are the final processing stages that relay signals to the mOFC. This does not mean that the other areas, especially the ones in this study in which a particular pattern emerged, do not contribute to the output from the OFA and the FFA to the mOFC; that output may depend upon earlier dialogue between “face-perceptive” areas which have been posited to have widespread connections amongst themselves (Elbich et al., 2019). It is possible that activity from various other “face perceptive” areas, especially the cuneus and the pSTS, in which a pattern emerged during the experience of facial beauty, is only communicated to the FFA and the OFA when a significant configuration in the activity emerges the former. Given that activity in mOFC correlates with activity in “face perceptive” areas only when a particular pattern of activity emerges in the latter, our results suggest that the anatomical outputs from “face-perceptive” areas to FFA and OFA and from the latter to mOFC are engaged only on a “need-to-know” basis. At present this must remain speculative until we have a better knowledge of the anatomical connections between these areas and until the chronology of activity between them is determined by future studies with much finer temporal resolution.

To the best current approximation, we can therefore conclude that the experience of facial beauty is determined by the emergence of a distinct pattern of activity in FFA, OFA and the parallel emergence of activity in the mOFC. By parallel we do not imply simultaneous; it stands to reason to suppose that activity in the FFA and the OFA precedes that in the mOFC. Our methods were ill suited to determine the chronology of activation, an issue that future studies will address.

## Conclusion

While there remain many details to settle, our study represents a first step in trying to understand the neural determinants of beauty. Our results lead us to conjecture that, through evolution, there has been a selection for beautiful faces and that this selection is represented neurally in specific patterns, or “significant configurations”, within specific brain areas that are strongly implicated in the perception of faces. The precise features that render a face beautiful, beyond the accepted general properties of symmetry, proportions and precise relationships – which are not in themselves necessarily sufficient to render a face beautiful – may be unknown. But we suggest that, whatever they may turn out to be, they are represented neurally by distinctive patterns of activity in the relevant sensory areas, and it is only when this pattern emerges that there is, as a correlate, activity within the mOFC. We conjecture that it may be the combined activity within these face perceptive areas and the mOFC that leads, as a correlate, to the experience of beauty.

## Methods

### Participants

19 healthy, right-handed adults (9 male, mean age 26.1 ± 1.6) participated in the study; of these, 8 were West Europeans, 1 was East European, 4 were Chinese, 3 Indian, and 3 Southeast Asian (Malaysia and Vietnam); none was an artist or had been trained in art history. Prior to the experiment. All participants gave written informed consent, and the experiment was approved by the ethical committee of Ludwig-Maximilians Universität Munich (LMU), in accordance with the principles of the Declaration of Helsinki. Two participants (Germans, 1 male and 1 female) were excluded because technical problems led to the loss of their data, leaving 17 complete datasets for analysis.

### Stimuli

The stimulus set consisted of 120 photographs of faces from our own database, supplemented by faces from the Chicago Face Data Base and the Face Research Lab London set. Based on the ratings given to the faces in these datasets, 40 faces (20 male) were selected to represent each of the three conditions - the “very beautiful”, “average”, and “not beautiful” conditions, respectively. The stimuli were standardised colour photographs, thus limiting the variation in low-level features such as brightness, which are unrelated to differences in individual faces (see Supplementary materials). All stimuli had the following characteristics: eye-gaze forward, head position forwards, and as neutral an expression as possible. We did not crop or otherwise modify the photographs to eliminate hair and ears, as is common; such manipulation gives the photographs an artificial mask-like appearance which is out of place in a study of facial beauty because what is removed commonly constitutes an important element in assigning an aesthetic status to a face; it is sometimes even not easy to determine the gender of such cropped faces with any certainty. All these features are ones, we believe, that subjects may take into consideration when rating faces according to beauty. Stimuli were back-projected onto a screen mounted in the scanner and viewed through an angled mirror on the head coil. Stimulus presentation was controlled by a personal computer running the ‘‘Presentation’’ software package (Neurobehavioral Systems, Inc., Albany, CA); images were scaled to 500 × 500 pixels and subtended approximately 10° ×10° of visual angle.

### Procedure

Subjects viewed the face stimuli in the scanner and rated them after viewing them according to how beautiful they found them to be on a scale of 1 to 7 (with 1 being “not beautiful” at all and 7 “very beautiful”) while the activity in their brains was being scanned (see Figure 1). Pressing one button on a customized pad increased the value while pressing another one decreased it, from the neutral setting of 4. Subjects attended a single 1 hr session and, before entering the scanner, were given a short practice run of 10 trials to familiarize them with the task. Each subject completed 5 experimental scanning sessions consisting of 24 trials each. Each scan contained 8 faces from each of three conditions (“very beautiful”, “average”, “not beautiful”) and each face was shown only once. The order of presentation was counterbalanced across participants, and image order within each scan was pseudo-randomized and counterbalanced. At the end of the scanning experiment, subjects viewed and re-rated all the stimuli again, randomly and on the same scale of 1-7, according to how beautiful and how attractive they were, as well as how familiar the faces were to them.

The sequence within a trial was as follows: a blank screen with a central blinking fixation cross (1 sec) → stimulus appearance for 2 s → a blank screen for 5 sec a rating scale for 4.5 s during which subjects were asked to rate the face aesthetically. To avoid any confounds associated with the use of hand movement, the lower and higher values were counterbalanced across participants. After a 5 s inter-trial-interval, the next sequence was presented (Figure 1). Each scan also included 15 s of blank screen at the beginning and end of the cycle, to allow for better baseline signal estimation and removal of T1 saturation effects.

### Image acquisition

Brain images were acquired during daytime at the University Hospital of the LMU, using a 3.0 T system (Achieva, Philips Healthcare, Best, The Netherlands). For BOLD signals, T2*-weighted Echo Planar Imaging (EPI) sequences were used (repetition time (TR) = 2500 ms; echo time (TE) = 30 ms; ?ip angle = 90°; ascending acquisition, with an acquisition matrix of 80 × 80 and a slice thickness of 3 mm with no gap between slices). Each of the 5 functional runs included 180 whole brain acquisitions for each subject. Structural data was acquired with a T1-weighted scan of each participant’s brain anatomy (1 mm × 1 mm × 1 mm; 240 × 240 matrix, field-of view = 220 mm).

## Data analyses

### Preprocessing

Data were pre-processed using SPM12 software (http://www.fil.ion.ucl.ac.uk/spm/). Functional images were realigned, slice-time corrected and normalized to the MNI template (ICBM 152) with a 3 mm isotropic voxel size. The registration was done by matching the whole of an individual’s T1 image to the template T1 image (ICBM152), followed by estimating nonlinear deformations. The affine and nonlinear transformations were then combined to re-slice all the functional volumes into the MNI template. Data was spatially smoothed with a Gaussian kernel (full width, half maximum = 8 mm) for univariate analyses but not for the MVPC/RSA performance.

#### A. Univariate analyses

##### 1. Categorical analysis

For this analysis we used, out of 120 stimuli, 20 trials with stimuli given the highest rating (“very beautiful”), and 20 given the lowest ratings (“not beautiful”) to ensure only trials of interest (which had ratings of 1, 2, 6, 7) were included. We fitted a standard general linear model (GLM) to each subject’s data. Two regressors specified the onsets of the “very-beautiful” and the “not-beautiful” conditions. The duration of the regressors corresponded to the 2 s that the face was shown for, together with the ensuing blank screen (5 s) in the given trial. Head movement parameters calculated from the re-alignment pre-processing step were included as regressors of no interest. We used SPM’s canonical haemodynamic response function (HRF) to convolve the task-related regressors and the default high-pass filter of 128s. The resulting individual contrast images were used for random effect analysis at the group level. The average BOLD response across the brain produced by viewing “very-beautiful” faces was compared to that produced by viewing “not beautiful” faces; a reverse comparison (“not beautiful” *vs* “very beautiful”) was also conducted. For these *t*-tests, significant voxels initially passed a voxel-wise statistical threshold of *p* ≤ .0001, and a cluster-level threshold was obtained at the family-wise-error (FWE)-corrected statistical significance level of *p* < .05.

##### 2. Parametric analysis

The entire set of stimuli was used here to identify those brain regions in which activity changed linearly with increasing and/or decreasing beauty ratings. A GLM incorporating a single task effect (face presentation), a parametric regressor (indicating subjects’ ratings of each face) and nuisance regressors (head movement parameters derived from realignment corrections) was used to compute parameter estimates and *t* contrast images for each subject. In this way, the height of the expected HRF was parametrically adjusted for all face events as a function of each participant’s beauty ratings for each face. Brain regions that responded to the ratings parametrically were identified by contrasting the parametrically modulated predictors against baseline. The resulting individual contrasts from the first-level model were entered into one-sample t-tests for group analyses. All statistical parametric maps and statistics reported in the tables were thresholded at a voxelwise level of *p* < .0001, with FEW-corrected *p* < 0.05 at the cluster level.

##### 3. Psychophysiological Interaction Analysis (PPI)

In a PPI analysis, the time course of the activity in a specified seed region is used to model the activity in target brain regions. Through it, we detected brain regions showing significant connectivity with the seed region in the mOFC, during the “very beautiful” compared to the “not-beautiful” face conditions. The time course (the physiological variable) was extracted from the seed region in mOFC (MNI: x⍰=⍰-3, y⍰=⍰53, z⍰=⍰-4; 6⍰mm radius sphere at the local peak). The psychological variables in the analysis included the “very-beautiful” and the “not-beautiful” conditions and separate one-sample t-tests were conducted to detect the group-level functional connectivity patterns for the contrast between these two conditions.

#### B. Multivariate analyses

##### 1. Decoding analysis

Multivariate pattern analysis (MVPA) was used to test for finer-grained neural representations specific for “very-beautiful” and “not-beautiful” ratings, using Python packages (Nilearn^1^ and Scikit-learn^2^); it used the 10 trials with the highest ratings (out of 120) and the 10 with the lowest ones from each participant. Each stimulus presentation was modelled as a separate event and GLM betas were estimated using the canonical function in SPM12. The beta values contained in these maps allowed the construction of vectors that serve as inputs to the decoding algorithms. We obtained 20 beta maps per subject and only used those beta-series amplitude estimates for the following analyses:

###### 1.1 Searchlight analysis

Maps of classification performance were computed using a whole-brain “searchlight” approach (Kriegeskorte et al., 2006); this allows the mapping of local multivariate effects by sliding a spherical window over the whole brain and performing independent decoding analyses within each sphere. We used a “leave-one-subject-out (LOSO)” cross-validation scheme to predict whether the subject had viewed the “very-beautiful” or “not beautiful” faces during the trial corresponding to the activation pattern provided to the classifier. More specifically, in each iteration we employed a logistic regression classification algorithm using data from all but one subject and tested its performance in predicting the class label (i.e., whether it was “very-beautiful” or “not-beautiful”) of the test subject’s data. The searchlight decoding analysis was performed for a series of 4-mm-radius spheres moved throughout the volume of the brain and centred on each voxel while doing so. Maps of above-chance performance (decoding accuracy > 0.5) for the classifier were thus created.

###### 1.2 Statistical analysis

To determine the statistical significance of the MVPA analyses, we performed a permutation test (Nichols & Holmes, 2002); this assesses the significance of the average decoding accuracy at the group-level in a non-parametric manner. 5,000 iterations of the “very-beautiful”/ “not-beautiful” trial-wise labels were computed and used to generate a null distribution of the 2 summary statistic. We tested the hypothesis that classification performance was better than chance at a FEW-corrected p < 0.05.

###### 1.3. Confusion matrices within Regions of Interest (ROIs)

Our ROIs were defined by the “very-beautiful” vs “not beautiful” face classification results, as well as core face-processing regions including OFA, FFA and pSTS. Five spherical ROIs with an average radius of 9 mm were centred at or near the peak values of manually selected voxels. A confusion matrix was computed for each ROI by averaging probabilistic classifier predictions. The significance of classification performance across all participants was assessed through permutation testing (5000 iterations).

##### 2. Representational similarity analysis (RSA)

A beta-series GLM was used to extract an estimate of the response amplitude for each trial at each voxel in grey matter. The ratings of faces labelled “very-beautiful” (top 10 of 120 from each participant) and those labelled “not-beautiful” (bottom 10) were selected, and the corresponding beta images were submitted to an RSA using the NeuroRA^3^.

###### 2.1. Hypothetical models

The first step of the RSA was the modelling of predicted similarity matrices corresponding to the “very-beautiful” and the “not-beautiful” conditions. Two RSA models were examined; the first tested for responses in which the similarity was driven by the viewing of “very beautiful” faces. This model excluded the possibility that the similarity could be driven by responses to “not-beautiful” faces, by hypothesising that the representational patterns are not similar among “not-beautiful” faces (Figure 5C, Model 1). The second model examined responses in which the similarity was driven by “not-beautiful” faces. This model tested for regions in which the response to “not-beautiful” faces showed similar patterns, irrespective of the patterns produced by the viewing of “very beautiful” faces (Figure 5C, Model 2). These two matrices show hypothetical distances, with values of 1 expressing complete similarity between particular conditions, and of 0 expressing complete dissimilarity (see Figure 5C).

###### 2.2. Searchlight analysis

For information-based searchlight analyses, we used a 3 × 3 × 3 voxel searchlight. Within a given cube, the Euclidean distance between pairs of activity patterns was calculated, resulting in a representational dissimilarity matrix (RDM) at each searchlight location. The group’s RDM was estimated by averaging across all participants’ RDMs. Next, the brain RDMs were Spearman-rank correlated with the two models reflecting different hypotheses for the similarity structure of neural responses in the “very-beautiful” and “not-beautiful” conditions.

###### 2.3. Statistical assessment

The searchlight results formed a continuous statistical whole-brain map reflecting how well the hypothetical models fit in each of the local brain regions. The resulting correlation coefficients were converted to z-values using Fisher transformation, and submitted to a permutation test (5000 iterations) to identify voxels in which the correlation value was significant at p < 0.001.

###### 2.4. RDMs within ROIs

Additional follow-up RSA analyses were conducted within functionally defined ROIs. Six ROIs with an average radius of 9 mm were created including regions in mOFC, cuneus, FFA, OFA and pSTS; each was tested separately, by correlating our hypothetical model with brain RDMs with the 10 most and the 10 least beautiful face stimuli. The significance of the correlations was tested by a permutation test with randomized stimulus labels (5000 times for iteration).

## Acknowledgment

This work was supported by the Leverhume Trust, London.

We are especially grateful to Ernst Pöeppel for facilitating these experiments.

## Footnotes

1 http://nilearn.github.io/.

2 https://scikit-learn.org/stable/.

3 https://neurora.github.io/NeuroRA/.

