## Supplementary materials for "The Neural Determinants of Beauty" (from another version of this preprint: version 2, 2021-06-08)

| Value | Very beautiful | Average | Not beautiful |
| --- | --- | --- | --- |
| Mean | 195.84 | 199.28 | 173.53 |
| SD | 18.73 | 15.04 | 19.91 |

### Table 1. Brightness of the experimental stimuli

Note. Value represents the brightness of the colour. It ranges from 0 to 255, with 0 being completely dark and 255 being fully bright. SD refers to standard deviation.

|  | Very beautiful | | Average | | Not beautiful | |
| --- | --- | --- | --- | --- | --- | --- |
|  | Mean | SE | Mean | SE | Mean | SE |
| Beauty | 5.80 | 0.08 | 3.51 | 0.09 | 2.38 | 0.09 |
| Attractiveness | 5.34 | 0.06 | 2.86 | 0.06 | 1.93 | 0.05 |
| Familiarity | 3.55 | 0.08 | 2.88 | 0.07 | 2.66 | 0.07 |

**Table 2.** Evaluations (mean and standard error) in each dimension as a function of facial beauty.

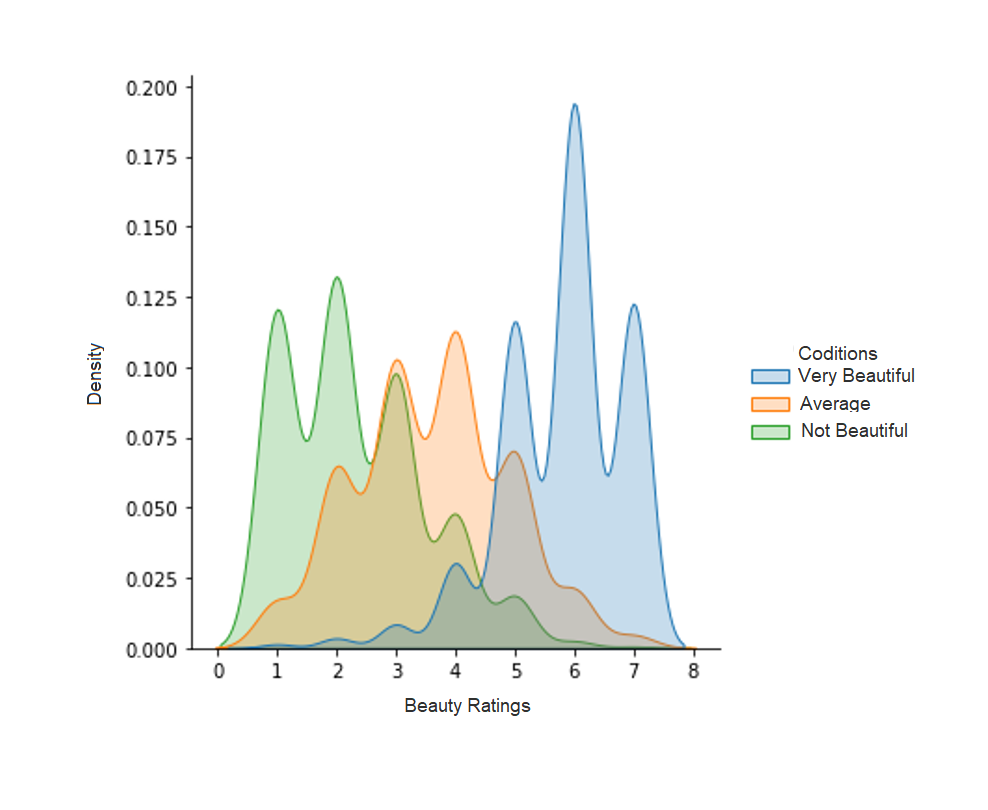

**Figure 1.** Density plot of beauty ratings in the experimental conditions. Blue indicates “very beautiful” faces, orange indicates “average” faces, and green indicates “not beautiful” faces.
